# Increased evolutionary rate in the 2014 West African Ebola outbreak is due to transient polymorphism and not positive selection

**DOI:** 10.1101/011429

**Authors:** Stephanie J. Spielman, Austin G. Meyer, Claus O. Wilke

## Abstract

Gire et al. (*Science* 345:1369–1372, 2014) analyzed 81 complete genomes sampled from the 2014 Zaire ebolavirus (EBOV) outbreak and reported “rapid accumulation of […] genetic variation” and a substitution rate that was “roughly twice as high within the 2014 outbreak as between outbreaks.” These findings have received widespread attention, and many have perceived Gire et al. (2014)’s results as implying rapid adaptation of EBOV to humans during the current outbreak. Here, we argue that, on the contrary, sequence divergence in EBOV is rather limited, and that the currently available data contain no robust signal of particularly rapid evolution or adaptation to humans. The doubled substitution rate can be attributed entirely to the application of a molecular-clock model to a population of sequences with minimal divergence and segregating polymorphisms. Our results highlight how subtle technical aspects of sophisticated evolutionary analysis methods may result in highly-publicized, misconstrued statements about an ongoing public health crisis.

---

Zaire ebolavirus (EBOV) is currently devastating West African populations in an unprecedented epidemic that has begun to spill over into many parts of the world. Since its discovery in the 1970s, EBOV has, at regular intervals, caused zoonotic outbreaks in human populations. Unlike past outbreaks, however, the current EBOV outbreak shows sustained transmission among humans, prompting concerns that as the outbreak escalates, EBOV may evolve to become endemic in humans. Recently, Gire et al. (2014) published 99 genomes from 78 patients infected in the current outbreak, sampled during May and June of 2014 in Sierra Leone. They analyzed 78 of these genomes (one from each patient), in combination with three EBOV genomes collected in Guinea during March 2014 (Baize et al., 2014), and reported “a rapid accumulation of…genetic variation.” They additionally stated that “[t]he observed substitution rate is roughly twice as high within the 2014 outbreak as between outbreaks.” The conclusions ultimately left readers, and indeed the scientific community at large, with the impression that EBOV is fast-evolving and possibly adapting to humans (see also Check Hayden 2014; Alexander et al. 2014).

By contrast, we do not find any robust evidence in the available 2014-outbreak EBOV genomes supporting this interpretation. While it is clear that mutations are certainly occurring in EBOV, the available genomic data do not show concrete evidence that EBOV is evolving particularly rapidly for an RNA virus. In fact, among the 81 genomes from the current outbreak, there are only 29 unique sequences (two from Guinea and 27 from Sierra Leone), and no genome contains more than 2 nonsynonymous mutations.

To put EBOV’s evolutionary dynamics into context, we compared the extent of genetic diversity within 2014-outbreak EBOV genes to the genetic diversity accumulated during the early months of the 2009 pandemic H1N1 influenza outbreak. Influenza virus is the archetypal rapidly-adapting human virus, and, like EBOV, it is a negative-sense single-stranded RNA virus. Since 1977, the two circulating seasonal influenza A strains have been H1N1 and H3N2. In late 2008 or early 2009, a new H1N1 strain, pandemic H1N1, a reassortant of several influenza viruses circulating in swine, was first transmitted from swine to humans and caused the 2009 H1N1 pandemic Smith et al. (2009). The first sample of pandemic H1N1 was identified on April 15, 2009 (Neumann et al., 2009). We considered here only H1N1 sequences collected in April 2009.

We constructed gene trees for EBOV nucleoprotein (*np*) and polymerase (*l*), selecting only genes from the 2014-outbreak EBOV sequences (Figure 1A). *np* and *l* have accumulated the most nucleotide sequence diversity of all seven EBOV genes during this outbreak, and thus they likely represent the most rapidly-evolving EBOV genes. We constrasted these phylogenies with gene trees for H1N1 hemagglutinin (*HA*) and nucleoprotein (*NP*) (Figure 1B) for sequences collected within a single month (April 2009) in the US. *HA*, the influenza surface protein responsible for host receptor-binding, is under intense selection pressure to evade host immunity, and it experiences elevated rates of nonsynonymous evolution near its receptor-binding site, though less so in H1N1 than in the more rapidly evolving H3N2 (Meyer et al., 2013). *NP* is not exposed on the viral envelope but evolves at a comparable rate to HA in pandemic H1N1 (Qu et al., 2011).

**Figure 1:**
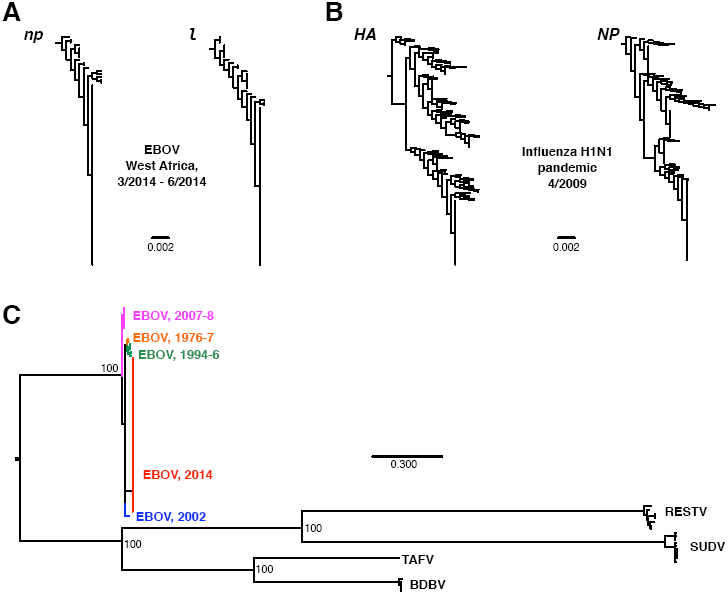
Limited divergence in EBOV 2014. Phylogenies in A and B were constructed from nucleotide data in FastTree2 (Price et al., 2009) under the GTR model. Sequences in A were restricted to 2014-outbreak EBOV sequences, and the phylogenies were rooted on *np* and *l* sequences sampled in 2002. Sequences in B were collected from the Influenza Research Database (Squires et al., 2012), with the search restricted to complete segments of pandemic H1N1 sampled during April 2009 in the US. Redundant strains, lab strains, and seasonal H1N1 were excluded. H1N1 phylogenies were rooted on the *HA* and *NP* genes from H1N1 strain A/Memphis/15/2000. The phylogeny in C was constructed from the genomic data of Gire et al. (2014) for all ebolavirus species, using RAxMLv.8.1.1 (Stamatakis, 2014) with the GTR+GAMMA model and a different partition for each gene. Numbers at nodes indicate bootstrap support. Abbreviations shown in the figure stand for Reston virus (RESTV), Bundibugyo virus (BDVD), Taï forest virus (TAFV), and Sudan virus (SUDV). All data and analysis scripts are available at https://github.com/wilkelab/EBOV_H1N1.

As the juxtaposed gene trees in Figures 1A and 1B demonstrate, even the fastest-evolving EBOV sequences from the 2014 outbreak are evolving much more slowly than H1N1 sequences do over similar temporal scale. Indeed, even the slowly-evolving influenza *NP* far outpaces EBOV genes in terms of accumulated genetic diversity. Furthermore, the average nucleotide diversities among the EBOV genes *np* and *l* from the current outbreak are approximately an order magnitude lower than the nucleotide diversities observed for the H1N1 genes *HA* and *NP*, while the mean root-to-tip distances are between 3 and 7 times lower in the EBOV genes than in the H1N1 genes (Table 1). Thus, EBOV is simply not accumulating mutations in a manner we would expect from a particularly rapidly evolving RNA virus.

**Table 1:** Mean root-to-tip distance and nucleotide diversity *π* for EBOV and influenza sequences. Nucleotide diversity was calculated as the average number of pairwise nucleotide differences among all sequences. All data and analysis scripts are available at https://github.com/wilkelab/EBOV_H1N1.

| **virus** | **outbreak** | **gene** | **mean root-to-tip distance** | **nucleotide diversity $\pi$** |
| --- | --- | --- | --- | --- |
| EBOV | 2014 | *np* | 0.0010 | 0.00025 |
|  |  | *l* | 0.0019 | 0.00016 |
|  | all (1976-2014) | *np* | 0.0078 | 0.0097 |
|  |  | *l* | 0.0080 | 0.0087 |
| H1N1 | April 2009 | *HA* | 0.0062 | 0.0018 |
|  |  | *NP* | 0.0055 | 0.0018 |

In fact, even considering all EBOV outbreaks since 1976, we find limited evidence for evolutionary divergence. For example, the mean root-to-tip distances in EBOV genes over all outbreaks (spanning four decades) are comparable to those found in influenza H1N1 virus in a single month (Table 1). Figure 1C shows a phylogeny of all EBOV sequences considered in Gire et al. (2014), as well as related ebolavirus species. The extent of sequence divergence across EBOV sequences collected from 1976 to present pales in comparison to divergence among ebolavirus species. Gire et al. (2014) reported a between-outbreak substitution rate of 0.8 × 10^−3^. By comparison, substitution rates in influenza range from 1.7 × 10^−3^ to 6 × 10^−3^, depending on the specific strain and gene considered (Rambaut et al., 2008; Smith et al., 2009; Bedford et al., 2010; Qu et al., 2011; Roche et al., 2014). The highest numbers, near 6 × 10^−3^, are observed in currently circulating H3N2 (Rambaut et al., 2008; Bedford et al., 2010).

The minimal sequence divergence within the current EBOV outbreak indicates that EBOV samples from the current outbreak should be considered to be drawn from a single, polymorphic population. Indeed, the mean pairwise sequence similarity among *unique* 2014-outbreak EBOV genomes is 99.84% (standard deviation of 0.39%). Moreover, according to Gire et al. (2014), there are at least 55 segregating mutations in the 2014-outbreak EBOV sequences, yet these mutations generally do not co-occur in any particular genome. For example, among these EBOV genome sequences, only three (two from Sierra Leone and one from Guinea) contain two nonsynonymous mutations, no genome contains three or more nonsynonymous changes, and there is no robust evidence that any given site has experienced multiple mutation events in the current outbreak.

Taken together, these results reveal that EBOV sequence data must be analyzed in a population genetics, rather than a purely phylogenetic, context. This requirement becomes particularly evident in substitution-rate estimates obtained under a molecular-clock model (as used in Gire et al. 2014). The molecular clock assumes that sequences have sufficiently diverged such that all differences are fixed substitutions rather than segregating polymorphisms. As a consequence, the rate of the molecular clock is highly time-dependent, such that the substitution rate is substantially elevated at short time-scales due to the confounding presence of segregating polymorphisms (Ho et al., 2005, 2007; Peterson and Masel, 2009; Ho et al., 2011). Gire et al. (2014) reported a doubling of the substitution rate in the current outbreak relative to a baseline EBOV substitution rate calculated from pooling all sequence data collected since 1976 (Figure 4F in Gire et al. 2014). While Gire et al. (2014) correctly stated that the doubled substitution rate they reported was “consistent with expectations from incomplete purifying selection,” the meaning of this short phrase is likely non-obvious to any scientist who is not an experienced evolutionary biologist. Indeed, other groups have simply quoted the doubled substitution rate without qualification regarding its cause (Alexander et al., 2014). We think it is important to emphasize that the doubled substitution rate is likely caused entirely by the short sampling time scale and contains no new information about Ebola biology in the current outbreak. Until more time has passed and mutations have either fixed or been removed from the population, results concerning EBOV substitution rate in the current outbreak are unreliable and inconclusive, and should clearly be labeled as such. For an alternative approach to analyzing EBOV adaption without focusing on evolutionary rate, see Łuksza et al. (2014).

In sum, we do not find any convincing evidence in the currently available 2014-outbreak EBOV data that this virus is particularly rapidly evolving or that non-synonymous mutations are accumulating. As more sequence data from the current outbreak are collected and made available for analysis, evidence may emerge to support such conclusions. However, until such data are released, we cannot deduce that EBOV 2014 sequences show signatures of increased evolutionary rate, much less of adaptation to humans.

## Acknowledgements

We would like to thank Trevor Bedford for helpful suggestions regarding our analysis of influenza data. This work was supported in part by DTRA grant HDTRA1-12-C-0007 and NSF Cooperative Agreement No. DBI-0939454 (BEACON Center). Computational resources were provided by the University of Texas at Austin’s Center for Computational Biology and Bioinformatics (CCBB).

## References

Alexander K A, Sanderson C E, Marathe M, Lewis B L, Rivers C M, Shaman J, Drake J M, Lofgren E, Dato V M, Eisenberg M C, Eubank S. 2014. What factors might have led to the emergence of Ebola in West Africa? PLOS Neglected Tropical Diseases, to appear. http://blogs.plos.org/speakingofmedicine/files/2014/11/Alexanderetal.pdf.

Baize S, Pannetier D, Oestereich L, Rieger T, Koivogui L, Magassouba N, Soropogui B, Sow M S, Keta S, De Clerck H, Tiffany A, Dominguez G, Loua M, Traor A, Koli M, Malano E R, Heleze E, Bocquin A, Mly S, Raoul H, Caro V, Cadar D, Gabriel M, Pahlmann M, Tappe D, Schmidt-Chanasit J, Impouma B, Diallo A K, Formenty P, Van Herp M, Gnther S. 2014. Emergence of zaire ebola virus disease in guinea. NEJM 371:1418–1425.

Bedford T, Cobey S, Beerli P, Pascual M. 2010. Global migration dynamics underlie evolution and persistence of human influenza A (H3N2). PLoS Pathog. 6:e1000918.

Check Hayden E. 2014. Ebola virus mutating rapidly as it spreads. Nature News, 28 August 2014. doi:10.1038/nature.2014.15777.

Gire S, Goba A, Andersen K, Sealfon R S, Park D, Kanneh L, Jalloh S, Momoh M, Fullah M, Dudas G, Wohl S, Moses L, Yozwiak N, Winnicki S, Matranga C, Malboeuf C, Qu J, Gladden A, Schaffner S, Yang X, Jiang P, Nekoui M, Colubri A, Coomber M, Fonnie M, Moigboi A, Gbakie M, Kamara F, Tucker V, Konuwa E, Saffa S, Sellu J, Jalloh A, Kovoma A, Koninga J, Mustapha I, Kargbo K, Foday M, Yillah M, Kanneh F, Robert W, Massally J L, Chapman S, Bochicchio J, Murphy C, Nusbaum C, Young S, Birren B, Grant D, Scheiffelin J, Lander E, Happi C, Gevao S, Gnirke A, Rambaut A, Garry R, Khan S, Sabeti P C. 2014. Genomic surveillance elucidates Ebola virus origin and transmission during the 2014 outbreak. Science 345:1369–1372.

Ho S Y, Phillips M J, Cooper A, Drummond A J. 2005. Time Dependency of Molecular Rate Estimates and Systematic Overestimation of Recent Divergence Times. Mol Biol Evol 22:1561–1568.

Ho S Y W, Lanfear R, Bromham L, Phillips M J, Soubrier J, Rodrigo A G, Cooper A. 2011. Time-dependent rates of molecular evolution. Mol Ecol 20:3087–3101.

Ho S Y W, Shapiro B, Phillips M J, Cooper A, Drummond A J. 2007. Evidence for time dependency of molecular rate estimates. Syst Biol 56:515–522.

Łuksza M, Bedford T, Lässig M. 2014. Epidemiological and evolutionary analysis of the 2014 Ebola virus outbreak. http://arxiv.org/abs/1411.1722.

Meyer A G, Dawson E T, Wilke C O. 2013. Cross-species comparison of site-specific evolutionary-rate variation in influenza hemagglutinin. Phil. Trans. R. Soc. B 368:20120334.

Neumann G, Noda T, Kawaoka Y. 2009. Emergence and pandemic potential of swine-origin H1N1 influenza virus. Nature 459:931–939.

Peterson G, Masel J. 2009. Quantitative prediction of molecular clock and Ka/Ks at short timescales. Mol Biol Evol 26:2595–2603.

Price M N, Dehal P S, Arkin A P. 2009. FastTree 2 – approximately maximum-likelihood trees for large alignments. PLOS ONE 5:e9490.

Qu Y, Zhang R, Cui P, Song G, Duan Z, Lei F. 2011. Evolutionary genomics of the pandemic 2009 H1N1 influenza viruses (pH1N 1v). Virology Journal 8:250.

Rambaut A, Pybus O G, Nelson M I, Viboud C, Taubenberger J K, Holmes E C. 2008. The genomic and epidemiological dynamics of human influenza A virus. Nature 453:615–619.

Roche B, Drake J M, Brown J, Stallknecht D E, Bedford T, Rohani P. 2014. Adaptive evolution and environmental durability jointly structure phylodynamic patterns in avian influenza viruses. PLoS Biol. 12:e1001931.

Smith G J, Vijaykrishna D, Bahl J, Lycett S J, Worobey M, Pybus O G, Ma S K, Cheung C L, Raghwani J, Bhatt S, Peiris J S, Guan Y, Rambaut A. 2009. Origins and evolutionary genomics of the 2009 swine-origin H1N1 influenza A epidemic. Nature 459:1122–1125.

Squires R, et al. 2012. Influenza research database: an integrated bioinformatics resource for influenza research and surveillance. Influenza Other Respir Viruses 6:404–416.

Stamatakis A. 2014. RAxML Version 8: A tool for phylogenetic analysis and post-analysis of large phylogenies. Bioinformatics 30:1312–1313.

